# The role of Anti-Mullerian hormone in predicting fertilization and pregnancy rates following in vitro fertilization-embryo transfer (IVF-ET) and intracytoplasmic sperm injection (ICSI) cycles at a public fertility centre in Nigeria

**DOI:** 10.1101/2020.09.07.285718

**Authors:** Safiyya Faruk Usman, Olubunmi Peter Ladipo, J.A.F Momoh, Chris Ovoroyeguono Agboghoroma, Nabila Datti Abubakar

## Abstract

**Objective:** To determine the role of Anti-Mullerian Hormone (AMH) in predicting fertilization and pregnancy rates following in vitro fertilization-embryo transfer (IVF-ET) and intracytoplasmic sperm injection (ICSI) treatment cycles.

**Methods:** This was a prospective cohort study of one hundred and fifty consecutive women undergoing IVF-ET/ICSI that were recruited from February 1, 2017 to October 31, 2018 at the Fertility centre of the National Hospital, Abuja, Nigeria. Participants’ plasma AMH were assayed and were followed up till achieving fertilization and pregnancy. Association between AMH levels, fertilization and pregnancy rates was assessed using univariable and multivariable logistic regression modelling to adjust for confounding variables.

**Results:** The mean age and mean AMH level of the participants were 36 ± 4.2 years and 1.74 ± 2.35ng/ml respectively. There was a statistically significant association between AMH level and age (P <0.001), duration of infertility (P =0.026), cause of infertility (P =0.035), number of oocytes retrieved (P =<0.001), number of embryos generated (P =<0.001) and type of treatment (P =<0.001). However, there was no significant difference in the fertilization rates (adjusted odds ratio [AdjOR] 0.36, 95% confidence interval [CI] 0.23–4.30; P =0.533) and pregnancy rates (AdjOR 0.26, 95% CI 0.04–2.00; P =0.210) at different plasma levels of AMH.

**Conclusion:** Plasma AMH level was not a predictor of fertilization and pregnancy rates among our cohort of patients who had IVF/ICSI treatment cycles.

## Introduction

As human fertility decreases globally, many couples may require assisted reproductive technology (ART).[1],[2] Counselling couples regarding their chances of a successful ART using an accurate prognostic test is necessary to obviate embarking on expensive treatment while minimal benefit is expected.[3] Considering the high cost, uncertainty of outcome and the possible complications of ART, exploring some parameters which could predict its outcome is of great value. Determinants of success in assisted reproduction are complex and a major factor in successful in-vitro fertilization (IVF) treatment is the ability of the ovary to respond to gonadotrophins stimulation and to develop multiple follicles. This response reflects the ovarian function or ovarian reserve (the functional potential of ovaries at any given time).[4] The ideal ovarian reserve test should aid identification of women with low chance of successful IVF consequent upon a reduced ovarian reserve. This will guide the decision concerning women to be excluded from further treatment and those requiring oocyte donation, so as effectively reduce costs of care for the couple and the health system.[5]

Although, age is an important determinant of ovarian response, there is a varying relationship between women’s reproductive capacity and chronological age.[1] With the paradigms of modern ART stressing the importance of treatment individualization and optimization, the need for more specific markers becomes essential.[6] Ovarian reserve can be assessed using endocrine markers such as Anti-Mullerian hormone (AMH), basal Follicle Stimulating Hormone (FSH), Inhibin B, Estradiol; sonographic examination of antral follicle count (AFC), ovarian volume and ovarian blood flow, and by ovarian stimulatory tests such as the clomiphene citrate challenge test (CCCT).^3^ The ultimate objective of these tests is to provide an accurate prediction of couples’ potential success prior to commencement of treatment, thus enabling a more feasible, patient-oriented treatment approach.[6] However, some endocrine markers are influenced by the menstrual cycle while inter- and intra-observer variation affects the accuracy of the ultra-sonographic markers.[7]

AMH, also called Mullerian inhibiting substance is a dimeric glycoprotein belonging to the transforming growth factor-β family.[8] It is secreted by the ovarian granulosa cells within the pre-antral and small antral follicles (<6mm in diameter). In the female fetus, production starts from as early as 36 weeks of gestation and continues until the menopause.[1],[9] AMH is increasingly recognised as superior to age, day-3 FSH, Estradiol or Inhibin B levels in predicting ovarian response.[5],[10],[11]

AMH has been demonstrated as being useful in individualising controlled ovarian stimulation to minimise treatment burden, reduce the risk of ovarian hyperstimulation syndrome and to maximise success rates.^4^ AMH level might thus inform individual women about their reproductive lifespan and current reproductive capacity.[10] Furthermore, some studies have also revealed significant positive correlation between AMH concentrations and pregnancy rate, ongoing pregnancy rate and live birth rate.[7],[2] However, results from some other reports indicated that the predictive value for serum AMH in relation to clinical pregnancy rate, ongoing pregnancy rate and live birth rate is controversial.[7] Consequently, the counseling and management of women with low AMH levels presents a significant challenge where either cycle cancellation or poor response is anticipated to avoid distress/disappointment.[4],[12]

The cost-effectiveness of the use of an AMH-based treatment strategy in IVF has recently been assessed, and proposed to lead to substantial savings.[13],[14] Furthermore, improving the success rate of IVF cycles will lessen the burden of infertility, as this is the procedure that has produced the highest pregnancy rate.[15] However, as ART is still relatively new in Nigeria, there is limited available data on the relationship between AMH and pregnancy rates of IVF cycles and the results have not been consistent across all studies.[3] Therefore, the aim of this study was to determine the role of AMH in predicting fertilization and pregnancy rates following IVF-ET/ICSI treatment cycles as the stratification of care based on AMH levels may optimize treatment outcomes.

## Materials and methods

A prospective cohort study was conducted among 150 consecutive consenting women with infertility who presented to the IVF Centre, National Hospital Abuja (NHA), Nigeria from February 1, 2017 to October 31, 2018, for IVF/ICSI treatment cycles. Women between the ages of 18-40 years with morphologic evidence of normal right and left ovaries on transvaginal ultrasound scan, normal menstrual cycle (24-35 days) and normal uterine cavity confirmed by previous hysteroscopy or hysterosalpingography were recruited. Women with characteristics that might affect reproductive outcome, such as previous history of ovarian surgery; endometriosis; endocrinological disorders (abnormal testosterone, abnormal prolactin, diabetes mellitus); hormonal therapy in the past 3 months; previous cancer chemotherapy; and male factor infertility were excluded from the study.

Socio-demographic, gynaecological, obstetric and past medical history of the participants was obtained using an interviewer-administered questionnaire. Further information was collected from the hospital records of the participants. About 5ml of blood was collected from the participants on day 2-5 of the menstrual cycle, prior to downregulation for AMH assay. Samples were immediately centrifuged to separate the plasma and stored in aliquots at −20°C. The samples were pooled and assayed at the same time to minimize intra-assay variation. Plasma levels of AMH was determined using Cobas e411^®^ auto analyzer (Roche, Basel, Switzerland). The patients were then classified based on their plasma level of AMH into negligible, reduced, normal and excessive responders. Quality assurance was ensured through proper sample collection, processing and storage. Analytical variables were controlled for to ensure precision and accuracy.

The IVF-ET/ICSI treatments were carried out using the standard protocol. Pituitary down-regulation was achieved with a GnRH agonist injection, given daily starting from the mid-luteal phase of the menstrual cycle. Controlled ovarian hyper stimulation was achieved with variable amounts of human menopausal gonadotrophin (HMG), (between 75-300IU) or recombinant FSH 150IU daily (Bharat Serums and Vaccines Ltd, Ambarnath, India). Treatment was monitored by serial transvaginal ultrasound scans and ovulation induction was achieved with 5000 - 10,000IU of hCG (Bharat Serums and Vaccines Ltd, Ambarnath, India), when at least two to three follicles have attained a diameter of between 18-22mm. Oocytes were retrieved 34-36 hours after hCG administration through transvaginal ultrasound guidance. The number of retrieved oocytes were recorded.

Gamete handling was done using flushing medium and the pre-equilibrated SAGE 1 culture medium (Origio, Måløv, Denmark), during oocyte washing, insemination and embryo culture. The oocytes number, morphology as well as their maturity were assessed and recorded. They were prepared and treated either by conventional IVF or ICSI depending on the quality of the sperm cells. Evidence of fertilization was checked for by the following day, which was indicated by the presence of 2 pronuclei and embryo transfers were done on day 3-5 using a Wallace Sure-Pro Ultra catheter® (Origio, Måløv, Denmark). Luteal phase support was achieved with intravaginal progesterone pessary Cyclogest® 400mg (Teva UK Ltd, Essex, England) per vaginum, twice daily and oral Oestradiol Valerate 2mg (Progynova®; Bayer Plc, Berkshire, UK) twice daily. The cycle was cancelled if day 9-10 folliculometry revealed one or no developing follicle, if no oocytes were retrieved, or if fertilization failed.

Serum β-hCG levels were assessed on the 14^th^ day post embryo-transfer and a positive test is interpreted as pregnancy. Clinical pregnancy was diagnosed by ultra-sonographic visualization of one or more gestational sacs two weeks after serum pregnancy test.[16] There was no case of ectopic pregnancy. For the purpose of this study, follow-up ended with a negative pregnancy test or the detection of clinical pregnancy after a positive pregnancy test.

The outcome measures were number of oocytes retrieved, number of embryos generated, fertilization rates (the number of fertilized eggs relative to the number of retrieved oocytes)[1], biochemical pregnancy rates (a pregnancy diagnosed only by the detection of β-hCG in serum or urine)[16] and clinical pregnancy rate (the number of clinical pregnancies per 100 initiated cycles).[16]

The study was approved by the Institutional Review Board (IRB) of the National Hospital, Abuja before initiation of the study protocol.

The information obtained from participants and the outcome were transferred from an excel spreadsheet to Stata 15.0 (Stata Corporation, College Station, Texas) statistical software for analyses. Frequency distributions of variables were generated and presented in tables and charts. Categorical variables such as fertilization and pregnancy rates were expressed as absolute numbers and percentages. For analysis, the plasma level of AMH was classified into four groups: AMH level of < 0.15ng/ml, 0.15-1.14ng/ml, 1.15-2.56ng/ml and >2.56ng/ml, considered as negligible, reduced, normal and excessive response respectively.

Continuous variables such as AMH level, age and BMI were described using mean and standard deviation (±SD) while duration of infertility was described using median and interquartile range (IQR) and the variables were subsequently categorised. Chi-square test (or Fishers Exact test) were used to assess the relationship between the socio-demographic and gynecological characteristics and the categories of AMH.

The association between continuous variables and the four groups of AMH was conducted using the oneway analysis of Variance or Kruskal Wallis test. Post hoc Bonferroni test was then conducted to determine where the difference lie. For the logistic regression modelling, >50% was considered high fertilization rate while ≤50% was considered low fertilization rate[17]. Univariable and multivariable logistic regression modelling was conducted to evaluate the relationship between AMH levels and achieving fertilization. Factors that had univariable P value<0.2 were used to build the multivariable model in a stepwise regression modelling to adjust for confounding and assess the role of AMH as a predictor of fertilization. Similar regression modelling was conducted for relationship between AMH levels and biochemical pregnancy. A P value <0.05 (95% confidence interval) was considered as statistically significant.

## Results

Of the 150 women that had IVF/ICSI treatments and were enrolled into the study, 80% (n=120/150) completed the study. The mean age of the participants was 36 (± 4.2) years with a range of 25 to 40 years and about 75% (n=112) of the women had tertiary level of education (Table 1). The median duration of infertility was 7 years (IQR; 1-20 years) and about 61% (n=92/150) had secondary infertility while 36% (n=54/150) of the women had ovarian factor as the main cause.

**Table 1.** Distribution of socio-demographic, reproductive and treatment characteristics of the participants

| Covariates | Frequency (n=150) | Percentage (%) |
| --- | --- | --- |
| **Age group (years) (mean: 36 (± 4.2) years)** |  |  |
| Under 34 | 46 | 31 |
| 35 and above | 104 | 69 |
| **Educational Status** |  |  |
| None | 5 | 3 |
| Primary | 11 | 7 |
| Secondary | 22 | 15 |
| Tertiary | 112 | 75 |
| **Body Mass Index (median:28, IQR:19-40kg/m<sup>2</sup>)** |  |  |
| Underweight | 2 | 1 |
| Normal weight | 42 | 28 |
| Overweight | 67 | 45 |
| Obese | 39 | 26 |
| **Parity (median:0, IQR 0-3)** |  |  |
| Nulliparous | 107 | 71 |
| Primiparous | 30 | 20 |
| Multiparous | 13 | 9 |
| **Duration of Infertility (median:7 years, IQR 1-20 years)** |  |  |
| Under 5 years | 53 | 35 |
| 5-10 years | 53 | 35 |
| 10 years and above | 44 | 30 |
| **Type of Infertility** |  |  |
| Primary | 58 | 39 |
| Secondary | 92 | 61 |
| **Cause of Infertility** |  |  |
| Cervico-uterine | 32 | 21 |
| Tubal | 39 | 8 |
| Ovarian | 54 | 36 |
| Unexplained | 39 | 26 |
| Others | 13 | 9 |
| **Treatment Type** |  |  |
| Cancelled | 30 | 20 |
| IVF | 76 | 51 |
| ICSI | 44 | 29 |
| **Protocol** |  |  |
| Long | 75 | 50 |
| Short | 75 | 50 |
| **Biochemical Pregnancy (Serum Beta HCG Level 14 days post Embryo-Transfer) \*** |  |  |
| <200ng/ml | 74 | 65 |
| ≥200ng/ml | 39 | 35 |
| **Clinical Pregnancy (Gestational Sacs at 6 weeks post Embryo-Transfer) \*\*** |  |  |
| 0 | 8 | 5.3 |
| 1 | 9 | 6 |
| 2 | 15 | 10 |
| 3 | 11 | 7 |
| 4 | 1 | 0.7 |

\*n=113, \*\*n=44, IQR: Interquartile range

Fifty-one percent (n= 76/150) of the women had IVF while 29% (n= 44/150) had ICSI. The treatment was cancelled in 20% (n=30/150) of the women which was all due to poor response. Half of the women had down-regulation through long protocol while the remaining half had short protocol. Thirty-nine participants achieved biochemical pregnancy while 36 achieved clinical pregnancy giving a biochemical pregnancy and clinical pregnancy rates of 26% (n=39/150) and 24% (n=36/150) respectively. Two patients (1.3%) developed ovarian hyperstimulation syndrome (OHSS).

The mean AMH level was 1.74 ± 2.35 ng/ml. The minimum level was 0.01ng/ml while the maximum was 12.8ng/ml. Seventy-eight participants had plasma AMH level of <0.15ng/ml (negligible response) while thirteen women had a normal response with plasma level of 1.15-2.56ng/ml (Fig 1). Seventy percent of the women (n=84/120) had good fertilization rate (>50%). The highest pregnancy rate of 58% (n=70/120) occurred within the group with the normal AMH level (Fig 2).

**Fig 1.**
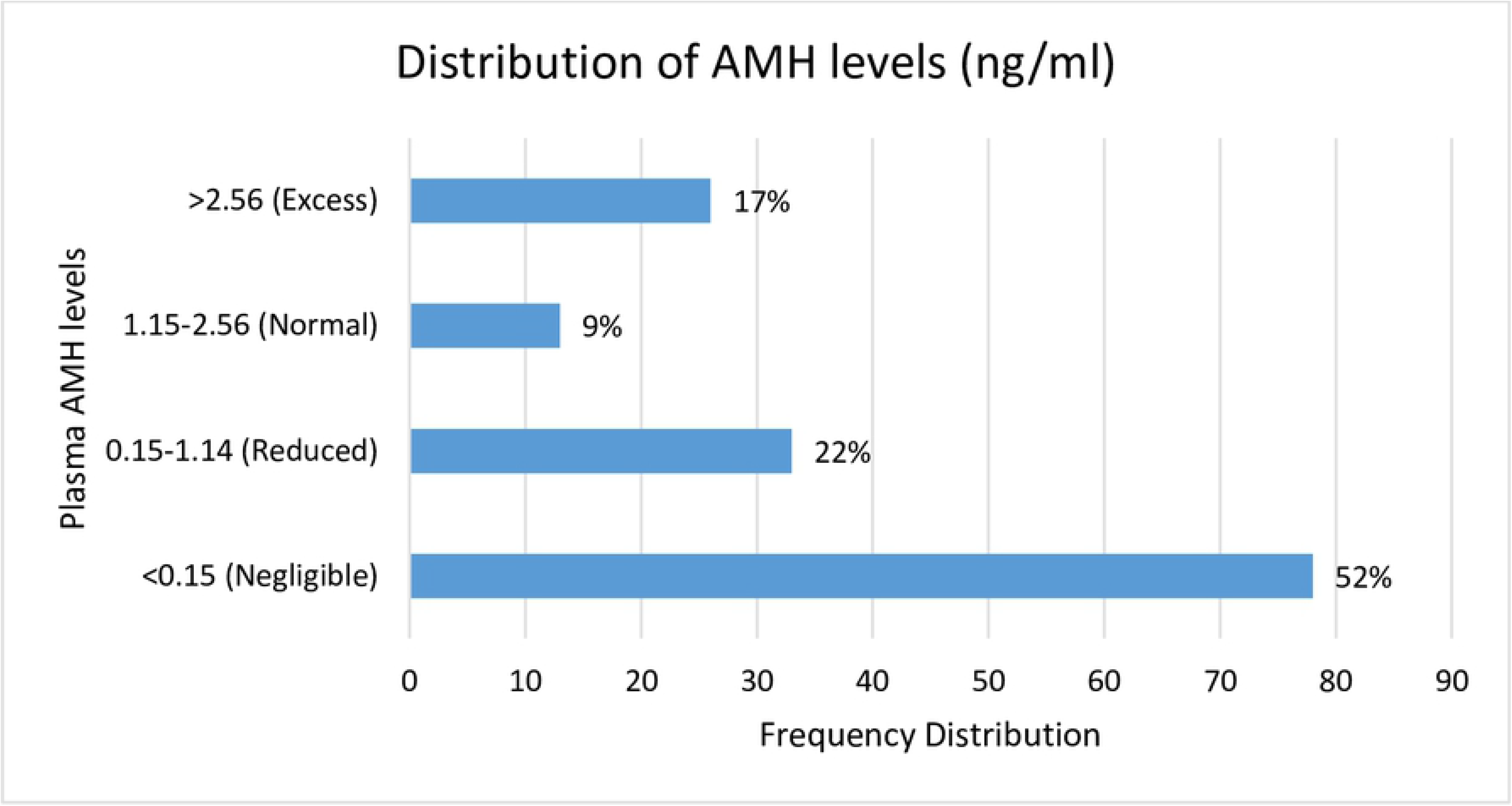
Frequency distribution of the plasma level of AMH of the participants.

**Fig 2.**
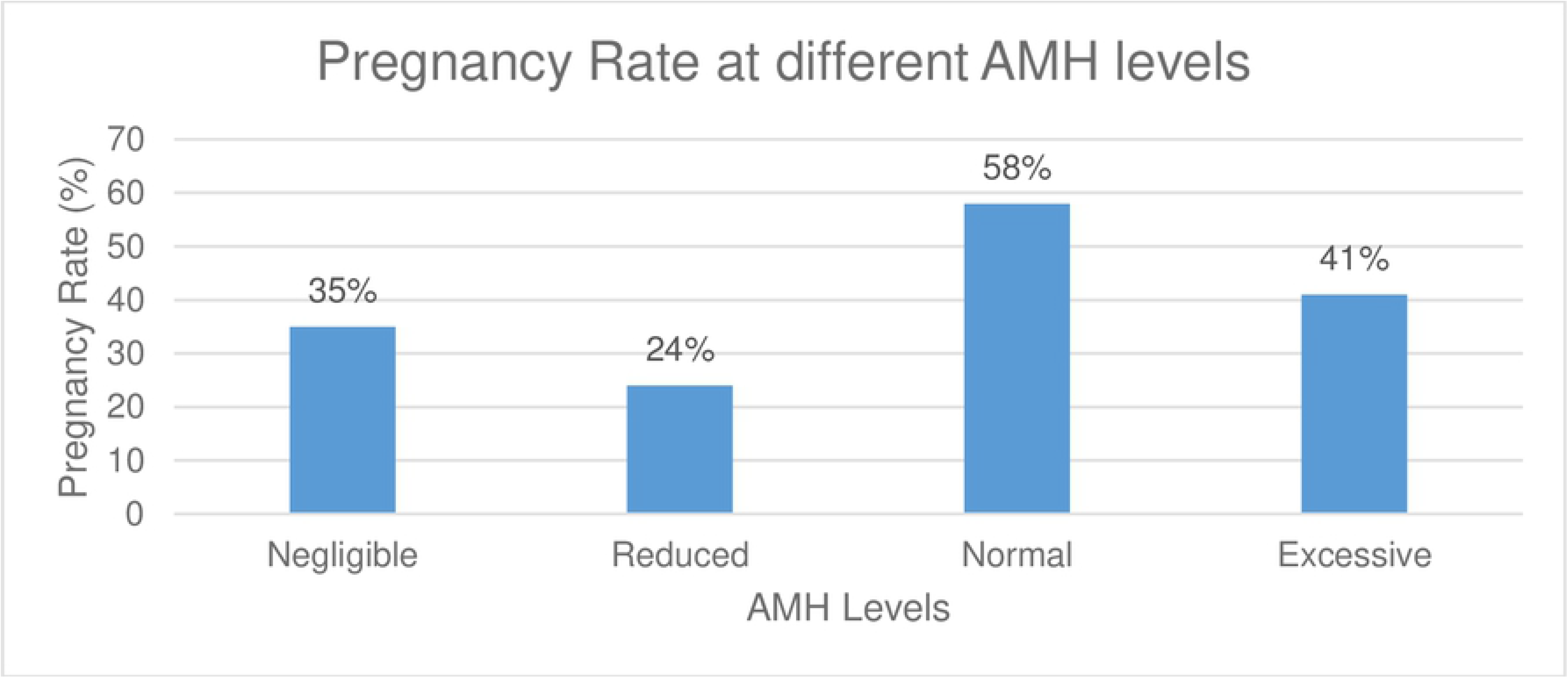
Pregnancy rate following IVF/ICSI treatment cycles among different plasma AMH levels.

There was a statistically significant difference in age (P value = 0.001), duration of infertility (P value = 0.026), cause of infertility (P value = 0.035), number of oocytes retrieved (P value = 0.001), number of embryos generated (P value = 0.001) and type of treatment (P value = 0.001) and the four groups of AMH levels. However, no differences were found among the four groups in terms of their BMI, parity, type of infertility and stimulation protocol used (Table 2).

**Table 2.** Association between AMH levels and different variables

| Covariates (n=150) | AMH Level (%) |  |  |  | Chi-square | P value |
| --- | --- | --- | --- | --- | --- | --- |
|  | Negligible (<0.15ng/ml) Frequency (%) | Reduced (0.15-1.14ng/ml) Frequency (%) | Normal (1.15-2.56ng/ml) Frequency (%) | Excessive (>2.56ng/ml) Frequency (%) |  |  |
| Age group (years) (Mean age ± SD) | 38 ± 3.4 | 36 ± 4.0 | 33 ± 4.9 | 34 ± 4.1 |  | 0.001 |
| Under 34 | 13 (17) | 10 (30) | 8 (62) | 15 (58) | 21.951 | 0.001 |
| 35 and above | 65 (83) | 23 (70) | 5 (38) | 11 (42) |  |  |
| Body Mass Index (Mean BMI ± SD) | 27 ± 4.5 | 28 ± 4.5 | 28 ± 5.0 | 27 ± 5.5 |  | 0.852 |
| Underweight | 2 (3) | 0 (0) | 0 (0) | 0 (0) | 9.805* | 0.367 |
| Normal weight | 18 (23) | 9 (27) | 4 (31) | 11 (42) |  |  |
| Overweight | 38 (49) | 12 (36) | 8 (62) | 9 (35) |  |  |
| Obese | 20 (26) | 12 (36) | 1 (8) | 6 (23) |  |  |
| Parity (Mean parity ± SD) | 0.5 ± 0.9 | 0.2 ± 0.6 | 0.3 ± 0.6 | 0.6 ± 1.1 |  | 0.353 |
| Nulliparous | 53 (53) | 27 (82) | 10 (77) | 17 (65) | 2.859* | 0.414 |
| Primiparous | 17 (17) | 5 (15) | 2 (15) | 6 (23) |  |  |
| Multiparous | 8 (8) | 1 (3) | 1 (8) | 3 (12) |  |  |
| Duration of Infertility (Mean Duration of Infertility ± SD) | 8.1 ± 4.9 | 6.8 ± 3.8 | 6.5 ± 3.9 | 6.9 ± 5.3 |  | 0.428 |
| Under 5 years | 22 (29) | 10 (30) | 6 (46) | 14 (54) | 14.36* | 0.026 |
| 5-10 years | 27 (35) | 18 (54) | 4 (31) | 4 (15) |  |  |
| 10 years and above | 28 (36) | 5 (15) | 3 (23) | 8 (31) |  |  |
| Type of Infertility |  |  |  |  |  |  |
| Primary | 28 (36) | 17 (52) | 4 (31) | 9 (35) | 3.071 | 0.381 |
| Secondary | 50 (64) | 16 (48) | 9 (69) | 17 (65) |  |  |
| Cause of Infertility |  |  |  |  |  |  |
| Cervico-uterine | 11 (14) | 12 (36) | 3 (23) | 6 (23) | 22.25* | 0.035 |
| Tubal | 19 (24) | 10 (30) | 5 (38) | 5 (19) |  |  |
| Ovarian | 37 (47) | 5 (15) | 3 (23) | 9 (34) |  |  |
| Unexplained | 4 (5) | 2 (6) | 2 (15) | 5 (19) |  |  |
| Others | 7 (9) | 4 (12) | 0 (0) | 1 (4) |  |  |
| Protocol |  |  |  |  |  |  |
| Long | 38 (49) | 18 (55) | 7 (54) | 12 (46) | 0.555 | 0.907 |
| Short | 40 (51) | 15 (45) | 6 (46) | 14 (54) |  |  |
| No. of Oocytes Retrieved (Mean No. of Oocytes Retrieved ± SD) | 3.6 ± 4.4 | 6.8 ± 5.4 | 8.1 ± 4.9 | 11.8 ± 7.0 | | 0.001 |
| 0 | 22 (28) | 5 (15) | 0 (0.00) | 2 (8) | 38.214* | 0.001 |
| 1-3 | 30 (38) | 5 (15) | 2 (15) | 2 (8) |  |  |
| 4-10 | 18 (23) | 16 (48) | 8 (62) | 10 (38) |  |  |
| >10 | 8 (10) | 7 (21) | 3 (23) | 12 (46) |  |  |
| No. of Embryos Generated (Mean No. of Embryos Generated ± SD) | 1.9 ± 0.7 | 2.3 ± 0.9 | 2.6 ± 0.8 | 2.8 ± 0.9 | | 0.001 |
| 0 | 26 (33) | 7 (21) | 0 (0.00) | 3 (12) | 34.239* | 0.001 |
| 1-3 | 38 (49) | 11 (33) | 7 (54) | 4 (15) |  |  |
| 4-10 | 13 (17) | 12 (36) | 4 (31) | 15 (58) |  |  |
| >10 | 1 (1) | 3 (9) | 2 (15) | 4 (15) |  |  |
| Treatment Type |  |  |  |  |  |  |
| Cancelled | 22 (28) | 6 (18) | 0 (0.00) | 2 (8) | 30.276* | 0.001 |
| IVF | 16 (21) | 10 (30) | 1 (8) | 17 (65) |  |  |
| ICSI | 40 (51) | 17 (52) | 12 (92) | 7 (27) |  |  |
| Serum β-HCG Level** (Mean Serum β-HCG Level ± SD) | 260.2 ± 738.4 | 134.2 ± 389.0 | 211.6 ± 292.7 | 226.1 ± 377.7 | | 0.853 |
| <200ng/ml | 35 (65) | 20 (80) | 6 (50) | 13 (59) | 4.012 | 0.260 |
| ≥200ng/ml | 19 (35) | 5 (20) | 6 (50) | 9 (41) | | |
| Gestational Sacs** (Mean gestational sacs ± SD) | 3 ± 2.7 | 3 ± 1.3 | 4 ± 1.7 | 3± 1.7 | | 0.047 |
| 0 | 3 (6) | 3 (12) | 1 (8) | 1 (4) | 18.365* | 0.244 |
| 1 | 7 (13) | 0 (0) | 0 (0) | 2 (9) |  |  |
| 2 | 6 (11) | 1 (4) | 4 (33) | 4 (18) |  |  |
| 3 | 4 (7) | 3 (12) | 2 (17) | 2 (9) |  |  |
| 4 | 0 (0) | 0 (0) | 0 (0) | 1 (5) |  |  |

\*Fisher's Exact Test, \*\* n=113, SD: Standard Deviation

The ANOVA test showed that there was a statistically significant difference in the mean age across the four groups of AMH. Mean age was highest among women with negligible AMH level (38 ± 3.4 years) and lowest in women with normal AMH level (33 ± 4.9 years), P value = 0.001 (Table 2). The post-hoc test showed that there was statistically significant difference between the mean age of women with normal versus negligible AMH levels (33 ± 4.9 Vs 38 ± 3.4 years, P value = 0.004) and between women with excessive versus negligible AMH levels (34 ± 4.1 Vs 38 ± 3.4, P value = 0.001).

There was also statistically significant difference in the mean number of oocytes retrieved across the four groups of AMH. The mean number of oocytes retrieved was lowest among the women with negligible AMH level (3.6 ± 4.4 oocytes) followed by reduced AMH level (6.8 ± 5.4 oocytes), normal (8.1 ± 4.9 oocytes) and then excessive AMH level (11.8 ± 7.0 oocytes), P value = 0.001 (Table 2). The post-hoc test showed that there was difference between the mean number of oocytes retrieved of women with reduced versus negligible AMH levels (6.8 ± 5.4 Vs 3.6 ±4.4 oocytes, P value = 0.020), between women with normal versus negligible AMH levels (8.1 ± 4.9 Vs 3.6 ± 4.4 oocytes, P value = 0.026) and between women with excessive versus negligible AMH levels (11.8 ± 7.0 Vs 3.6 ± 4.4 oocytes, P <0.001).

There was no statistically significant difference in the odds of achieving fertilization among the women with the different AMH categories (unadjusted odds ratio [UOR] 0.58, 95% confidence interval [CI] 0.09-3.36, P = 0.488). This relationship persisted after adjusting for the effect of age, BMI, duration of infertility, type of infertility, treatment protocol, number of oocytes retrieved, number of embryos generated, number of embryos transferred and type of treatment (AdjOR 0.36, 95% CI 0.23-4.30, P = 0.533) (Table 3).

**Table 3.** Logistic regression analysis showing the crude and adjusted odd ratios of AMH predicting fertilization and associated factors among the study participants

| Covariates | Crude (Unadjusted) |  |  | Adjusted* |  |  |
| --- | --- | --- | --- | --- | --- | --- |
|  | Odds Ratios | 95% CI | P value | Odds Ratios | 95% CI | P value |
| AMH Level |  |  |  |  |  |  |
| Negligible | 0.381 | 0.07-1.88 | 0.488 | 0.276 | 0.03-2.25 | 0.533 |
| Reduced | 0.314 | 0.06-1.68 |  | 0.187 | 0.02-1.82 |  |
| Normal | 1.000 |  |  | 1.000 |  |  |
| Excessive | 0.576 | 0.09-3.36 |  | 0.356 | 0.23-4.30 |  |
| Age group (years) |  |  |  |  |  |  |
| Under 34 | 1.000 |  | 0.437 | 1.000 |  | 0.486 |
| 35 and above | 0.703 | 0.29-1.70 |  | 1.605 | 0.42-6.07 |  |
| Body Mass Index |  |  |  |  |  |  |
| Underweight | 1 |  | 0.983 | 1 |  | 0.686 |
| Normal weight | 1.000 |  |  | 1.857 | 0.42-8.16 |  |
| Overweight | 1.081 | 0.44-2.64 |  | 1.000 |  |  |
| Obese | 1.083 | 0.38-3.11 |  | 1.144 | 0.25-5.24 |  |
| Duration of Infertility |  |  |  |  |  |  |
| Under 5 years | 1.000 |  | 0.578 | 1.000 |  | 0.972 |
| 5-10 years | 0.809 | 0.30-2.14 |  | 0.986 | 0.20-4.74 |  |
| 10 years and above | 0.594 | 0.22-1.60 |  | 0.842 | 0.17-4.06 |  |
| Type of Infertility |  |  |  |  |  |  |
| Primary | 1.000 |  | 0.728 | 1.000 |  | 0.113 |
| Secondary | 1.868 | 0.39-1.92 |  | 1.345 | 0.10-1.28 |  |
| Treatment Protocol |  |  |  |  |  |  |
| Long | 1.000 |  | 0.049 | 1.000 |  | 0.469 |
| Short | 0.457 | 0.21-0.99 |  | 0.666 | 0.22-1.99 |  |
| No. of Oocytes Retrieved |  |  |  |  |  |  |
| 0 | 1 |  | 0.169 | 1 |  | 0.002 |
| 1-3 | 2.911 | 0.94-8.97 |  | 29.50 | 4.26-204.49 |  |
| 4-10 | 1.000 |  |  | 1.000 |  |  |
| >10 | 1.129 | 0.40-3.18 |  | 0.644 | 0.15-2.81 |  |
| No. of Embryos Generated |  |  |  |  |  |  |
| 0-3 | 1.000 |  | 0.001 | 1 |  | 0.001 |
| >3 | 6.257 | 2.22-17.71 |  | 15.65 | 2.93-83.49 |  |
| Treatment Type |  |  |  |  |  |  |
| IVF | 1.000 |  | 0.453 | 1.000 |  | 0.834 |
| ICSI | 1.429 | 0.56-3.63 |  | 1.138 | 0.34-3.82 |  |

\*Adjusted for age, BMI, duration of infertility, type of infertility, treatment protocol, number of oocytes retrieved, number of embryos generated and type of treatment

Similarly, there was no significant difference in the odds of achieving pregnancy among women with different categories of AMH (UOR 0.49, 95% CI 0.11-2.06, P = 0.244). This relationship also persisted even after adjusting for the effect of age, BMI, duration of infertility, type of infertility, treatment protocol, number of oocytes retrieved, number of embryos generated, number of embryos transferred, type of treatment and fertilization rate (AdjOR 0.27, 95% CI 0.04-2.00, P = 0.210) (Table 4).

**Table 4.**
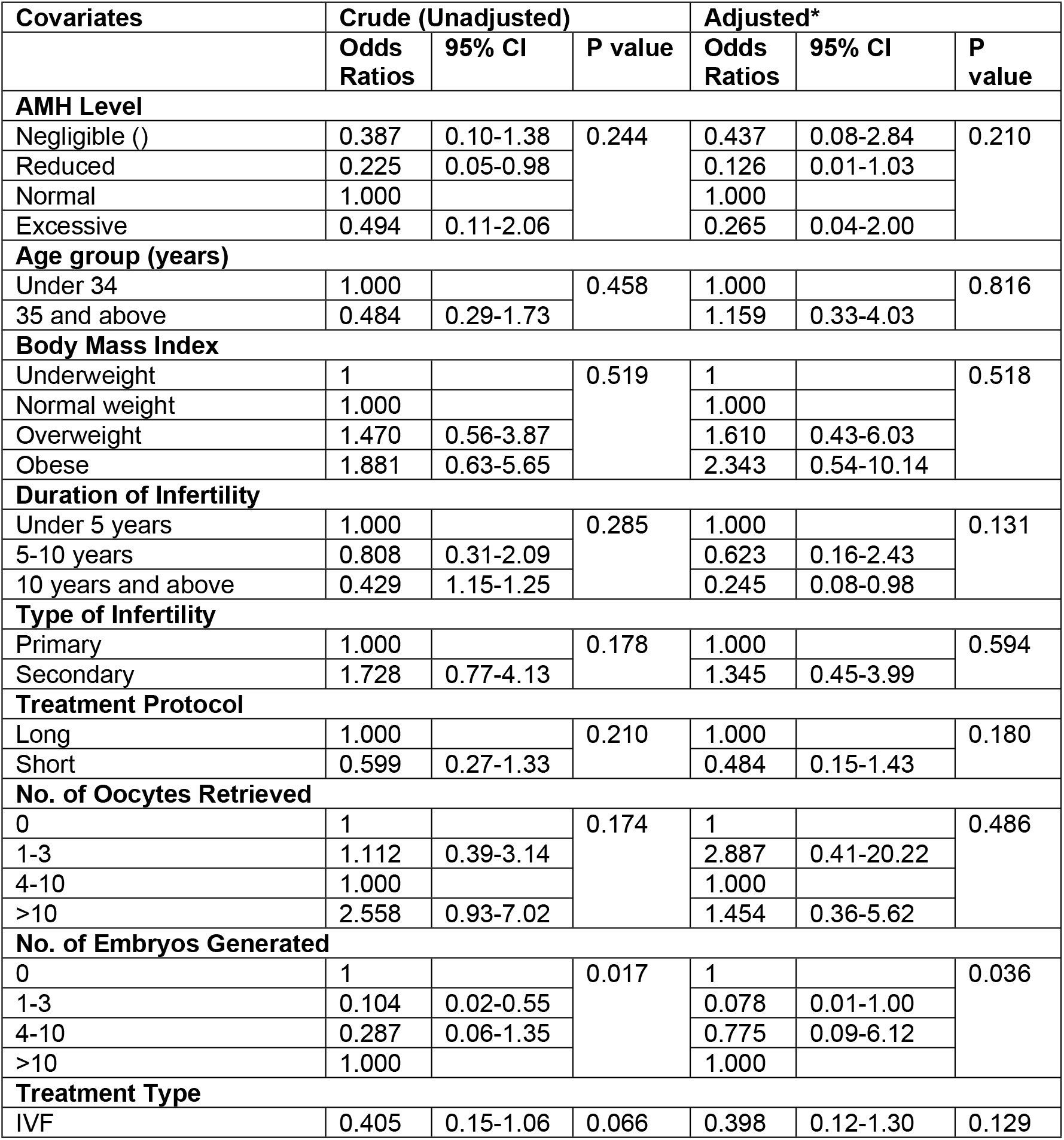
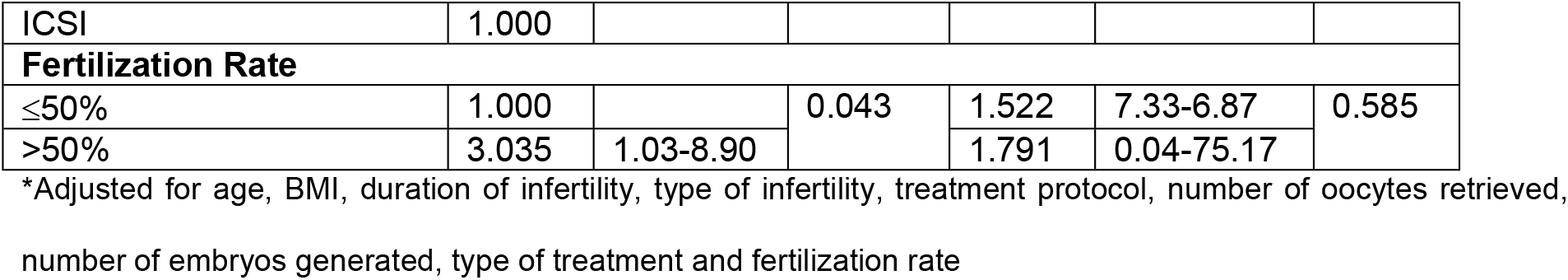
Logistic regression analysis showing the crude and adjusted odd ratios of AMH predicting pregnancy and associated factors among the study participants

## Discussion

In this study, the plasma AMH concentration of the majority (78%) of the women that had IVF/ICSI was found to be negligible (<0.15ng/ml). This might be due to the advanced age at presentation, as AMH decreases with advancing age. More so, ART is usually the last resort in most resource-poor countries like Nigeria because of poor availability and accessibility.[18] The result of this study suggests that there was significant association between plasma AMH concentration and age. AMH levels was found to fall with increasing age and this is consistent with findings from existing literature.[19] There was no significant difference found between AMH and BMI which was similar to a previous study that revealed that changes in AMH may be explained only by changes in age, as BMI significantly increased with ageing.[19] Similarly, there was no significant difference found between AMH and parity, which was consistent with a study that showed that pregnancies and number of offspring are distributed in an AMH unrelated pattern.[19]

The statistically significant difference found between AMH levels and number of oocytes retrieved was similar to findings by Rong Li et al where serum AMH concentration was positively correlated with the number of oocytes retrieved in a cohort of Chinese infertile women.[20] The higher the level of AMH, the higher the oocyte yield, which was similar to findings reported by Kevin Keane et al and Scott Nelson et al where AMH was found to be strongly correlated with oocyte yield.[21],[22] The number of oocytes retrieved has been recognised to affect the outcome of an IVF/ICSI cycle.[21] Hence, low levels of AMH is a marker of either cycle cancellation or poor response to ovarian stimulation. In this study, out of the 30 women that had cycle cancellation, 73% had negligible ovarian response (AMH level <0.15ng/ml). The association found between AMH and the number of embryos generated was also similar to findings from previous studies.[8],[23]

The multivariable logistic regression analysis demonstrates that there was no significant difference in fertilization rate and pregnancy rate among the four groups of AMH level even after adjusting for the effect of other variables. This suggests that AMH level has not been shown to predict fertilization and pregnancy rates following IVF/ICSI treatments, despite being able to demonstrate response to ovarian hyperstimulation. This is consistent with other studies where serum levels of AMH were not significantly associated with fertilization rates[7],[2],[8] and pregnancy rates.[8],[24],[25],[26] This finding might be attributable to the fact that though oocyte number and quality decline with age, fertility varies significantly even among women of the same age.[27] Further explanation can be derived from a study by Norbert Gleicher et al which found that at varying peripheral serum concentrations, AMH, demonstrates hitherto unknown and contradictory effects on IVF outcomes.[27] Additionally, a retrospective study by Nigel Pereira et al found that in patients with diminished ovarian reserve who have good quality embryos, AMH is not associated with clinical pregnancy, spontaneous miscarriage or live birth rates.[28]

On the contrary, some studies have revealed significant positive correlation between AMH concentrations and pregnancy rate and ongoing pregnancy rate.[7],[2],[21] Even though these studies use similar IVF protocols, they were however, large and retrospective.

To our knowledge, this is the first study addressing the relationship between AMH and fertilization and pregnancy rates in sub-Saharan Africa and, specifically, Nigeria. Other strengths of this study were the availability of a reputable IVF centre where facility-related and procedure-related adverse effects on IVF/ICSI outcomes are unlikely. The study was the first of its kind in my centre, thereby providing the background for further research in the field. Additionally, the study population was clearly outlined and confounding variables were controlled for in the analysis. The use of a fully automated, fast, sensitive and highly precise method of AMH measurement was another strength of this study.

The limitations of this study include the skewing of the participants to the older age range as most patients for IVF do not present early in this environment. This in turn might be responsible for some form of sampling bias. Furthermore, although this study has presented a detailed analysis of the relationship between AMH and fertilization and pregnancy rates, it was constrained by the non-availability of genetic screening of embryos to rule out the effect of genetic disorders on fertilization and pregnancy rates.

Nonetheless, the study adds to the limited body of literature regarding AMH as a predictor of IVF outcomes and would be of interest to experts involved with fertility treatments especially during counselling of women prior to IVF/ICSI on the role of AMH on the prognostication of outcome. In addition to AMH, an important predictive factor for IVF success is age, further studies may consider evaluating the role of AMH on IVF/ICSI treatment outcomes in women over 40 years.

